# Spatiotemporal cell landscape of human embryonic tooth development

**DOI:** 10.1101/2023.03.01.530693

**Authors:** Yueqi Shi, Yejia Yu, Jutang Li, Shoufu Sun, Li Han, Shaoyi Wang, Ke Guo, Jingang Yang, Wenjia Wei, Jin Qiu

## Abstract

Understanding the cellular composition and trajectory of human tooth development is valuable for dentistry and stem cell engineering research. Previous single-cell studies have focused on mature human tooth and developing mice tooth, but cell landscape on human embryonic dental development is still lacking. We collected tooth germ tissues from aborted fetus (17-24 week) for single cell RNA sequence and spatial transcriptome. We classified all cells into seven subclusters of epithelium, seven clusters of mesenchyme and other cell types like Schwann cell precursor and pericyte. For epithelium, the Stratum intermedium branch and the ameloblast branch diverged from a same set of KRT15+-HOPX+-ALCAM+ epithelial stem cell lineage, but the spatial distribution of two branches were not clearly distinct. This trajectory received spatially adjacent regulation signals from mesenchyme and pericyte, including JAG1 and APP. The differentiation of pulp cell and pre-odontoblast showed four waves of temporally distinct gene expression, which involved regulation networks of LHX9, DLX5 and SP7 and were regulated by upstream ligands like BMP family. Taken together, we provided a reference landscape for the research on human early tooth development, covering different spatial structures and developmental periods.

## Introduction

Human tooth development is a long and complex process starts at embryonic stage and last until adolescent period. This process represents a good model of organogenesis involving epithelial-mesenchyme interaction, morphogenesis and adult tissue self-renewal[1]. At the embryonic stage, dental development started at the formation of mesenchyme from neural crest[2] and the dental placode originated from the thickened oral epithelium. At this stage, dental epithelium starts to proliferate in some regions of the dental lamina, which then migrates to the deep connective tissue, and the end-stage epithelium proliferate and further differentiate to form the enamel organ. In parallel to this process, the ectomesenchyme layers beneath the proliferative epithelium also proliferate rapidly and aggregate around the epithelial layer to form the condensed mesenchyme layer. The epithelia and mesenchyme that proliferate in this restricted regions together consist in the tooth germ[3], which is the structural basis of further dental development The tooth germ is made up of three parts: 1. Oral ectoderm-originated enamel organ, which give rise to enamel; 2. Ectomesenchyme-originated dental papilla tissue, which give rise to dentin and pulp tissue; 3. Ectomesenchyme-originated dental follicle, which give rise to cementum, periodontal ligament and alveolar bone[4]. A comprehensive understanding of dental development would require a cell atlas that cover all these heterogenous cell types and tissues.

The rapid development of single cell techniques provides new opportunity for this task[5]. Recent single-cell studies on mouse dental tissues have elucidated the cell heterogeneity from early state Cranial neural crest cells[6] to later incisor tissues[7, 8], and delineated the differentiation and migration trajectory of epithelial and mesenchymal cells. Other studies using dental tissues from healthy volunteers also constructed dell landscape of postnatal human dental pulp[9, 10], apical papilla[7], periodontal tissues[9, 11] and immature tooth germ[12]. Based on these and other existing data, Hermans et al.[13] built a comprehensive dental cell landscape that basically covered all developmental stages of mouse incisor and molar tissue as well as human postnatal dental tissues, which has been proved valuable for clinical and bioengineering studies. However, some challenges remain unsolved by these existing efforts. An obvious blank is the lack of data on human prenatal dental tissues. Since mouse incisor has everlasting self-renewal capacity and exhibits dramatic physiological difference compared with human[14], the immediate translation of dental development knowledge from mouse to human should be interpreted with caution. Another drawback is the insufficient spatial resolution: current single-cell analysis relied on immunofluorescence that tags one or two proteins to study the spatial distribution of cell subtypes, which inevitably missed more refined information on developmental process based on multiple genes.

To fill in these blanks, we collected primary and permanent tooth germ tissues from 17 to 24 gestational week aborted fetus and applied single-cell RNA sequencing (scRNA) and spatial transcriptome (ST) to construct a spatiotemporal cell landscape of early human tooth development. Our study aimed to address the following questions. First, the cellular composition of human embryonic tooth germ at different stages. Second, the developmental trajectory of dental epithelium and mesenchyme, as well as their underlying biological process, gene regulatory networks and intercellular signaling pathways. Third, the spatial distribution of these elements.

## Method

### Sample collection and preprocessing

This study is approved and supervised by the ethic committee of Shanghai Tongren Hospital. Five fresh human fetal tooth germ tissues were collected from elective, nonmedically motivated pregnancy terminations at different embryonic developmental stages (2, 2 and 1 sample at 17, 20 and 24 PCW, respectively), without evidence of developmental abnormalities. Informed consent was obtained from each donor before termination of pregnancy. The parents of the fetus we chose decided to terminate the pregnancy for personal reasons. All prenatal examinations showed that the fetus had no developmental abnormalities. The method of pregnancy termination was medically induced. The parents of the fetus are healthy, with no developmental abnormalities, and there is no family history of facial and dental developmental abnormalities. Details of sample preprocess was provided in Supplementary methods.

### scRNA and ST

Library construction, sequencing and data filtration sere described in the Supplementary methods. Both libraries of scRNA and ST were sequenced with paired-end 150 bp sequencing (PE150) by NovaSeq 6000 platform. We retained non-doublet cells with number of detected genes >500 and <2500, and mitochondria gene percentage <5%. Library size normalization was performed with NormalizeData function in Seurat[15] to obtain the normalized count. Specifically, the global-scaling normalization method “LogNormalize” normalized the gene expression measurements for each cell by the total expression, multiplied by a scaling factor (10,000 by default), and transformed by Seurat: SCTransform function. Cell phase was defined by Seurat: CellCycleScoring function based on cell cycle gene list[16]. Spatial transcriptome data were preprocessed by Seurat in a similar way.

### Cluster analysis

For scRNA data, we applied Louvain community detection-based cluster analysis implemented in Seurat: FindClusters function with number of principle component=20 and resolution=0.5. We applied Seurat: FindMarkers Wilcoxon test to find top 15 markers for each cluster, and defined them according to existing knowledge on dental cell types. Epithelium and mesenchyme cell were further extracted and underwent second round of clustering, with the same parameters.

### Pseudotime analysis

For epithelium and mesenchyme (excluding isolated follicle and MSC clusters) cell groups, we applied monocle3:learn_graph and order_cells function[17] to learn the pseudotime trajectory. We then applied monocle3: graph_test function on top 5,000 variable genes detected by Seurat: FindVariableGenes function to find all genes that significantly altered along the trajectory. Genes with FDR-adjusted p<0.01 were further used for module detection by monocle3: find_gene_modules function. For the ease of interpretation, in epithelium cell group we separately applied module detection in each branch, and only retained gene modules that showed up-regulation or down-regulation along each of the three branches.

### Transcription factor regulation network analysis

We applied SCENIC[18] to construct transcription factors (TF)-target genes regulation network, separately for epithelium and mesenchyme. TF-motif annotation was downloaded from SCENIC website, which was corresponded to hg38 refseq gene annotation and contained a 10kb window of each gene. We retained target genes for extended network of each TF, and applied Fisher test to see if module genes detected by pseudotime analysis were enriched in any of the TF regulation network. TFs with FDR-adjusted p<0.01 and Odds ratio >2 were retained for each module. We further extracted the overall expression of a TF and all its targets (so-called regulon) of each cell to verify that this expression value had the similar expression pattern as module genes.

### Gene Ontology analysis

We used ClusterProfiler R package[19] to apply Gene Ontology-Biological Process (GO-BP) analysis by a hypergeometry test. The background gene list was set as all genes with GO-BP annotations. Only terms with >5 and <500 genes were considered. We applied simplify function to remove highly similar terms. Only terms with FDR-adjusted p<0.01 were retained. The odds ratio and p value were reported in the word cloud plot.

### nichenetr analysis

We used nichenetr R package[20] to apply signaling network analysis as described previously[12]. We separately considered 1) up-regulation modules of each of the three epithelium branches; 2) immature (first and second wave) and mature (fourth wave) of mesenchyme differentiation as target gene list, and the final signaling network were combined in each of these two categories. Ligands were prioritized by area under curve (AUC) and Pearson correlation coefficient calculated by nichenetr.

### Spatial transcriptome analysis

After defining cell clusters on scRNA data, we projected it to spatial transcriptome data by Seurat: FindTransferAnchors and TransferData functions to estimate the cell type probability of each spot. We assign cell types with the highest probability to each spot. We used misty R package[21] to estimate the spatial relationship among highlighted cell types, TFs and expression levels of ligands and target genes. We chose the “intra” mode in misty analysis and trained a linear model to predict each of the target values (cell probability or gene expression) by adjacent predicting values. Spatial adjacency was quantified by importance score in the prediction model.

## Result

### Overview of cell types in human embryonic tooth

We collected tooth germ tissues from aborted fetus spanning 17 to 24 gestational weeks (Figure 1A) for scRNA and ST. In scRNA, we obtained expression data for 11,218 single cells of nine general cell types that passed quality control (Figure 1B-D and Figure S1). As shown in Figure 1D and Figure S1, we classified these cell groups based on gene panel (Table S1) from existing cell atlas of mouse tooth[7, 8], adolescent tooth[9] and histology knowledges. We first defined cells with epithelial origin based on expression of dental epithelial marker KRT5, KRT14, FXYD3 and COL17A1 (Figure 1D and S1). Mesenchyme origin cells were characterized by their expression of EMILIN1 and MSX1. An isolated subgroup of mesenchyme origin cells was defined as apical pulp-like cell based on expression of SFRP2. Lastly, we defined mitotic cells, pericyte, endothelium, neural crest cells (NCC), macrophage and lymphocyte based on their specific markers CENPF, RGS5, CD34, FOXD3+SOX10, C1QA and PTPRC+CD52, respectively (Table S1). Across different gestational weeks, we observed difference in the proportion of epithelium and pericytes, but no difference in cell cycle distribution (Figure 1C). We additionally applied differential expression analysis between developmental time points for pericyte, and found that there was little difference with biological significance. This result indicated that pericyte did not show significant differentiation between the two time points, thus we mainly focused on dental epithelium and mesenchyme clusters.

**Figure 1.**
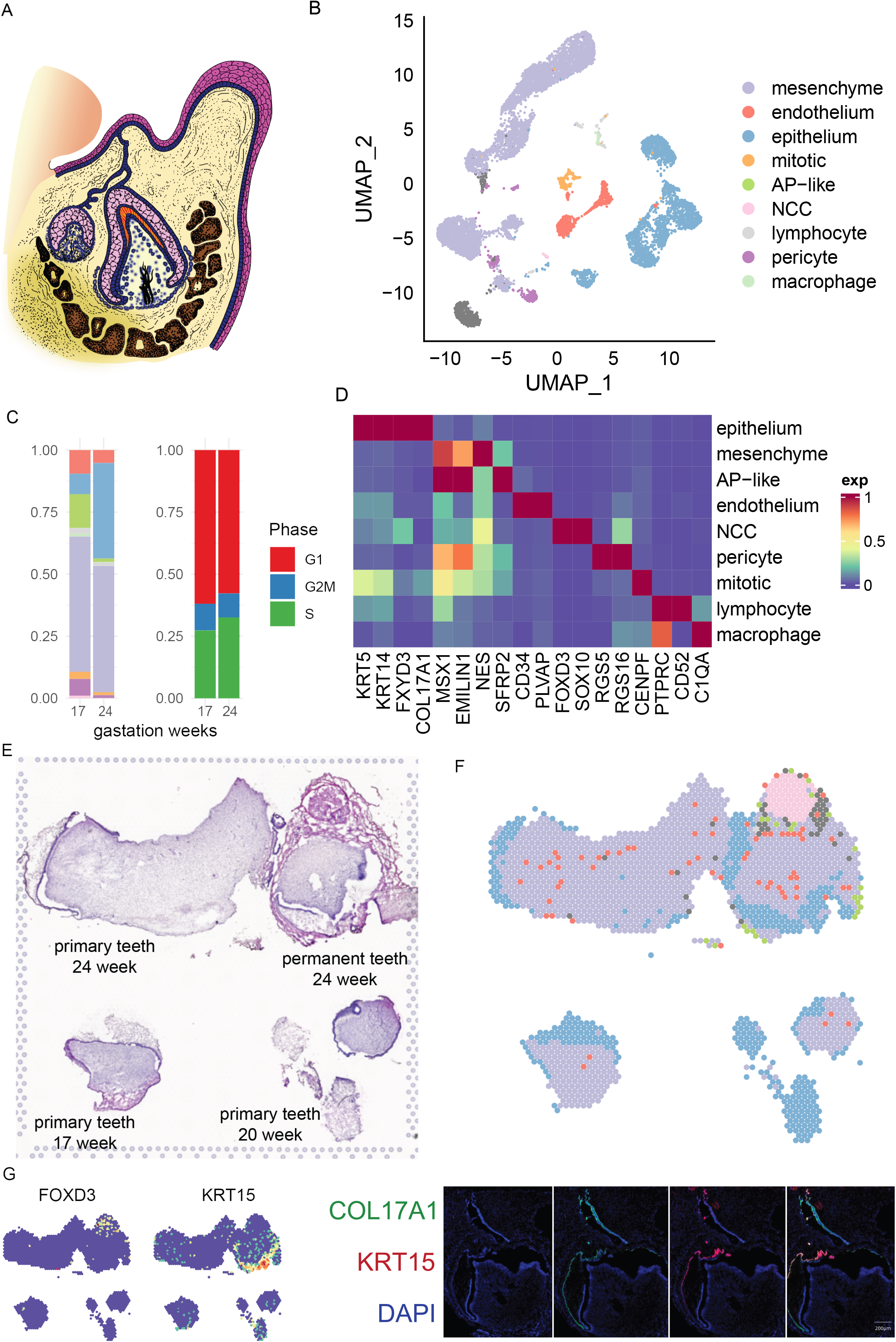
Overview of the human embryonic tooth cell atlas. A: schematic diagram of sample collection. B: UMAP plot of single-cell clustering. NCC: neural crest cell. AP: apical pappila. C: percentage of each cell type and cell cycle phase in samples from different gestational weeks. Cell cluster color corresponded to B. D: Marker scaled expression of each cluster. E: H-E staining of spatial transcriptome slice. F: Cell type annotation of each spot, defined as the cell type with the highest probability. G: Marker scaled expression of cell markers.

We further applied ST on tooth slices from three donors: a slice from 24-week embryo that contained both tooth germ of primary tooth and tooth germ of permanent tooth, a slice of 17-week primary tooth and a slice of 20-week primary tooth (Figure 1E). We applied anchor-based integration method implemented in Seurat v3[15] to evaluate the proportion of each cell cluster of scRNA in each spatial plot (Figure 1F and Figure S2). As expected, we observed that epithelium incompletely surrounded pulp tissues that were consisted mainly of mesenchyme cells and scattered pericytes and endothelium. AP-like cells were only observed in permanent tooth germ. We also found an isolated NCC group surrounded by endothelium, which were adjacent to permanent tooth germ and likely represented the process of neuron outgrowth. The spatial distribution of cell markers at transcription and protein levels also supported these spatial characteristics (Figure 1G). Taken together, we successfully constructed the cell type atlas of human embryonic tooth germ that covered various developmental timepoints and spatial structures.

### Branched trajectory of dental epithelium differentiation

Next, we conducted in-depth analysis of the cell constitution and development process of epithelium cell groups. By applying cluster analysis and pseudotime analysis, we partitioned all epithelial origin cells into three branches (Figure 2A). The root branch expressed GAS6 and IGFBP5, and consisted of two continuous cell clusters: outer enamel epithelium expressing KRT15 and SPARCL1 (OEE, Figure 2B), and inner enamel epithelium expressing CRABP1 and HOPX (IEE). Root branch ended at a cluster of ALCAM positive cells, which also expressed CD24 and HOPX which were associated with epithelial stem cells. ALCAM+ cell then gave rise to two branches: the SI branch consisted of SI progenitors expressing KRT7 and TAGLN and Stratum intermedium expressing CLDN10, and the ameloblast branch expressing ASCL5 and AMELX (Figure S3). Lastly, we found a small subcluster of ameloblast expressing SHH, WNT10A, and exclusively WNT10B, which we defined as enamel knot (EK). 24-week tooth germ contained moderately more ameloblast and less ALCAM+ SC than 17-week tooth germ, and the overall cell proportion was similar (Figure 2C). These results suggested that the development of SI and ameloblast have diverge trajectory, even though they share a same set of OEE-IEE lineage as progenitor. This result resembled previous identification of a dental progenitor that could give rise to both SI and ameloblast following injury[22], and suggested that the ALCAM+ population in our study might have similar roles as dental progenitors.

**Figure 2.**
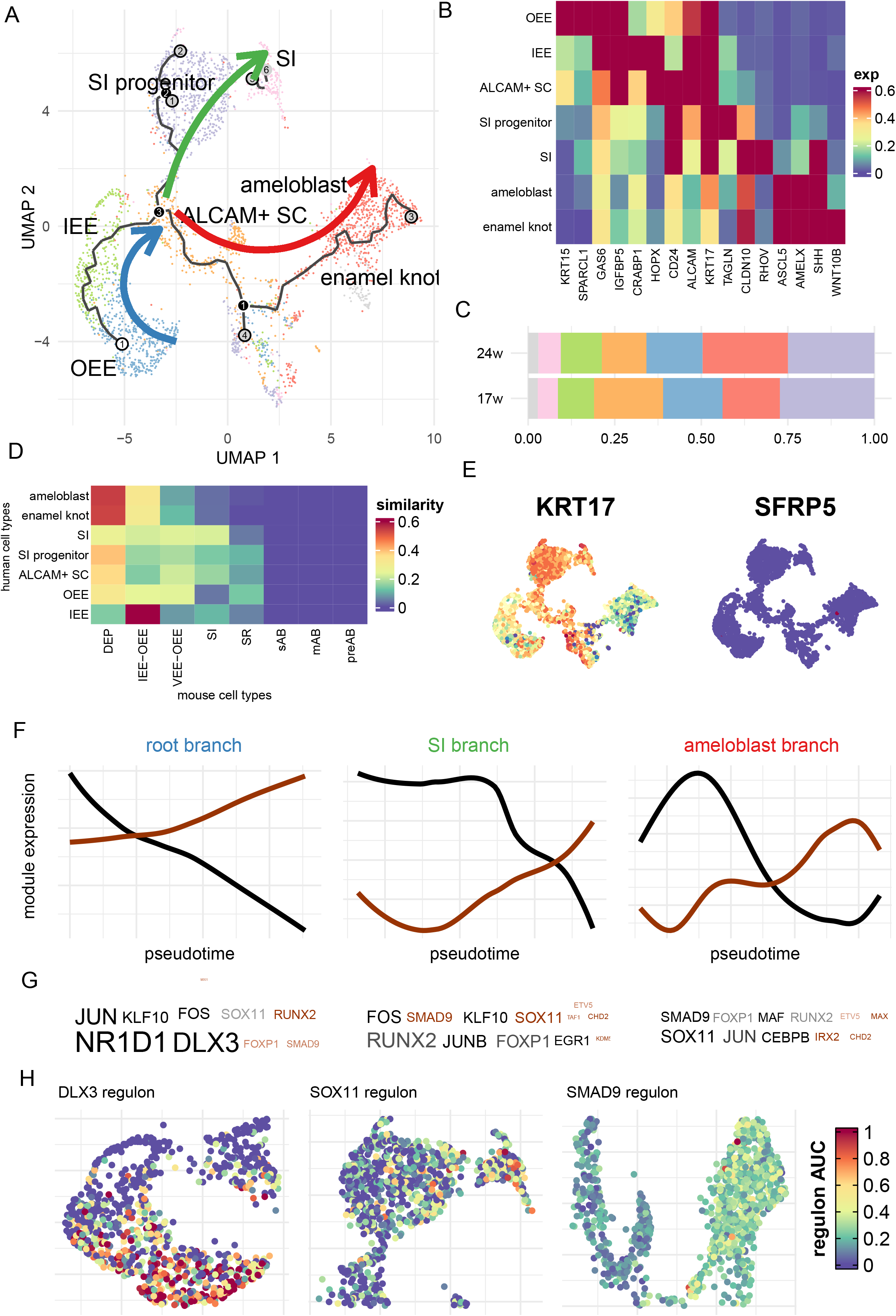
In-depth analysis of dental epithelium. A: UMAP plot and pseudotime analysis of epithelium cell clusters. Arrows marked development direction. OEE: outer enamel epithelium. IEE: inner enamel epithelium, EK: enamel knot. SC: epithelial stem cell. SI: Stratum intermedium. B-C: similar to Figure 1 C-D, but for epithelium clusters. D: Similarity between epithelial clusters from human and mouse embryonic tooth. DEP: dental epithelial progenitor. SR: stellate reticulum. sAB: secretary ameloblast. mAB: mature ameloblast. PreA: pre-ameloblast. E: expression level of human-specific SI marker KRT17 and mouse-specific DEP marker SFRP5. F: Module expression along pseudotime. For each branch, we identified an up-regulation module and a down-regulation module. G: Transcription factors whose targets were enriched for module genes. Color denoted the module that were enriched, transparency showed enrichment p-value, font size showed odds ratio. Only TF that showed non-zero expression were showed. H: Regulon expression of selected TF.

We further compared our human embryonic data with mouse data meta-analyzed by Hermans et al[13]. using anchored based integration. As shown in Figure 2D, human and mouse embryonic dental epithelium showed profound difference. Most of human cell types mostly resembled dental epithelium progenitor (DEP), and were dissimilar to their corresponding cell types in mouse except the IEE. We also found no similar cell types to any of the mouse ameloblast subtypes (pre-ameloblast, secretary ameloblast and mature ameloblast); in fact, human ameloblast and enamel knot were mostly similar to DEP in mouse. We also found considerable difference in cell marker expression. In human, KRT17 (Figure 2E) was found to be highly expressed in multiple epithelia especially the SI lineage, whereas in mouse there was no notable expression patterns. On the other hand, although most human epithelia was similar to mouse DEP, its marker SFRP5 (Figure 2E) showed almost zero expression in human dental epithelium.

We lastly analyzed the biological process underlying these branches. At the threshold of FDR-adjusted p<0.01, we found 622, 1397 and 842 genes whose expression levels significantly altered along pseudotime in root, SI and ameloblast branch, respectively. We applied gene module detection algorithm implemented in monocle3[17] to find genes that were consistently up-or down-regulated in each branch (Table S2-4). We ran SCENIC[18] to find transcription factors (TF) that regulate these genes. As shown in Figure 2F and Table S5-7, TF like NR1D1, DLX3, RUNX2, SOX11 and SMAD9 were predicted to regulate a large number of genes showing alterations in each branch. For example, among 18 target genes of DLX3, 14 were down-regulated in the root branch (Odds ratio, OR=102, Fisher test p=6.45×10^−18^). Consistently, the overall expression levels of DLX3 and its targets (so-called regulon[18]) were highly expressed in the primitive OEE, and gradually diminished in the root branch (Figure 2G). We highlighted regulons of several other TF showing consistent expression patterns in SI and ameloblast branch, such as SOX11 and SMAD9 that were activated during SI and ameloblast development (Figure 2G).

### Spatial patterns of dental epithelium development

We next linked the single cell-level signals to the spatial transcriptome. As shown in Figure 3A, B and Figure S4, OEE were only found in the outer surface of permanent tooth germ, whereas ALCAM+ SC were found inner to OEE in permanent tooth germ. Consistently, ameloblast were only sparsely distributed inner to stem cell layers in permanent tooth germ, but were abundant and densely distributed in primary tooth germ. Ameloblast were found at the interface between pulp tissue and epithelial tissues, whereas SI progenitor were distributed at the outer layers of ameloblast. These patterns were also supported by the spatial expression levels of the corresponding cell markers (Figure S5). We also summarized the inferred cell proportion of each spatial spot of all tissues (Figure 3C), and found that IEE-OEE were mainly distributed in permanent tooth germ, whereas Stratum intermedium were mainly found in tooth germ of 24 gestational week.

**Figure 3.**
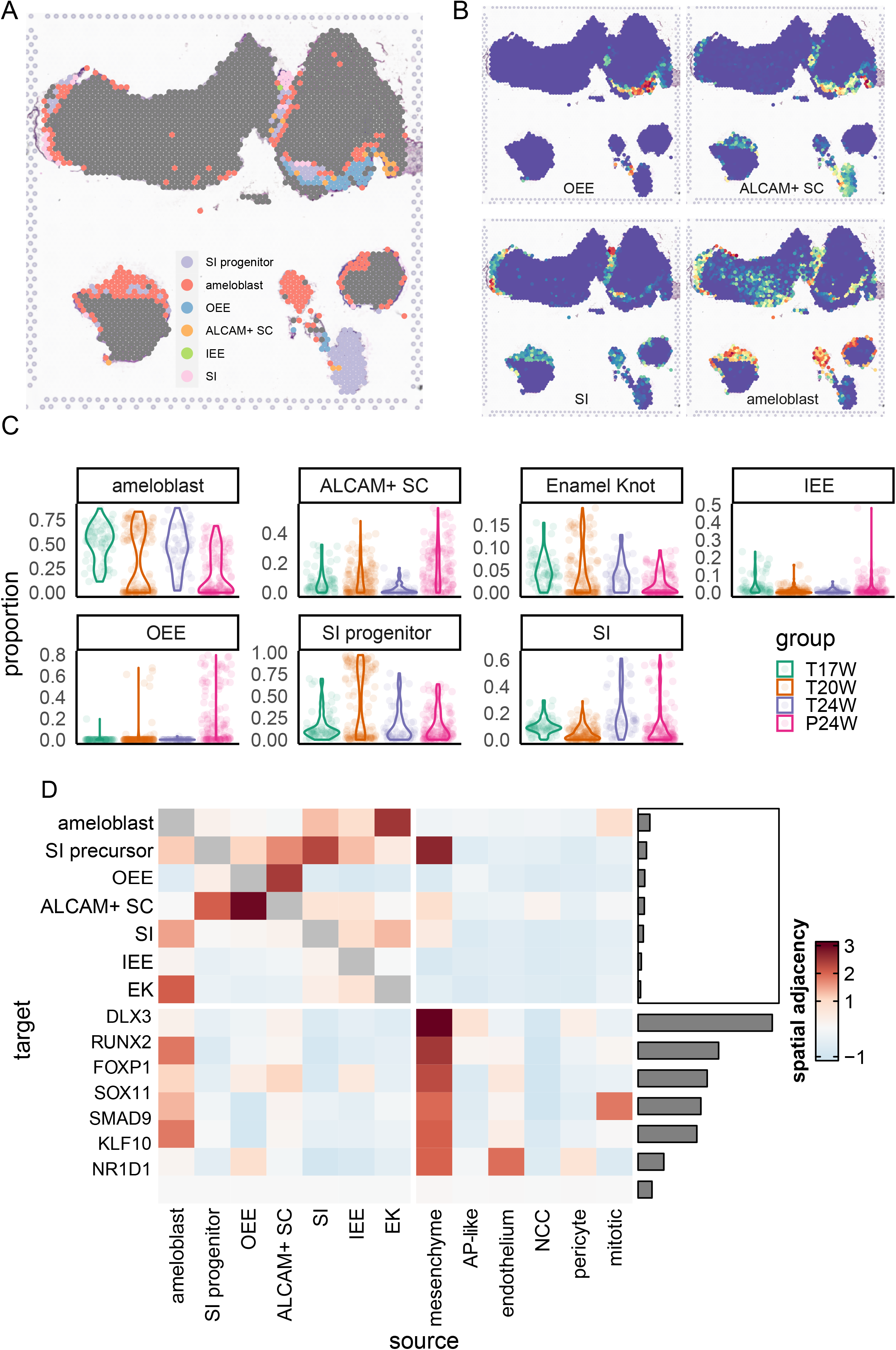
Spatial transcriptome of dental epithelium. A: Cell type annotation of each spot. B: Cell type probability spatial distribution. C: Cell type probability distribution per developmental stage. D: misty result of spatial adjacency, quantified by model importance score. Higher importance score indicated that the level of targets is determined by the adjacent level of source. RMSE: Root mean squared error of each model.

We then applied misty[21] to elucidate the spatial adjacency of different epithelial cell subtypes and key TFs. As shown in Figure 3D, when analyzing epithelial cells along, OEE and ALCAM+ SC had close distribution with each other, whereas ameloblast, enamel knot and Stratum intermedium aggregated together. Including all cell types together, only mesenchyme showed close relationship with multiple epithelial cells. When analyzing key TFs in epithelial differentiation (i.e., those TFs highlighted in Figure 2), we also observed that expression levels of various key epithelial TF was significantly associated to adjacent mesenchyme proportion (Figure 3D). In Conclusion, we inferred that signal from mesenchyme cells had an important role in regulating dental epithelial differentiation. This involved DLX3, which is known to be implicated in bone, hair and tooth development, and RUNX2, a key factor in osteoblast differentiation and skeletal morphogenesis, which were found notably modulated by mesenchymal signals. In fact, the intercellular communication between mesenchymal and epithelial cells appears to employ a complex signaling cascade, ensuring the temporal and spatial control of these molecules during the various stages of tooth formation.

### Involvement of LHX9, TCF7 and SP7 in dental pulp development

Similar to the analysis of dental epithelium, we further deciphered the spatiotemporal dynamics of mesenchyme cell group. As shown in Figure 4A-C and Figure S6, we identified seven subtypes of mesenchyme origin. Apical pulps (AP) were defined based on the expression of SFRP1, SOSTDC1 and SMOC2. We defined distal pulp according to expression of SOX9, which contained two subtypes: one isolated group expressed TNC, DKK3 and HEY1 (TNC+ DP) and another expressed FRZB, FGF3 and TWIST2 (TWIST2+ DP). We further defined two odontoblast groups by SALL1 expression, and distinguished them by FGF3+ TWIST2 expression (FGF3+ OD) and DKK3+FBN2 expression (FBN2+ OD). Lastly, we found two follicle cell clusters, one expressed IGFBP5+SPON1+FOXF1 (follicle 1) and another expressed GDF10 and COL12A1 (follicle 2). Compared with 24-week tooth germ, 17-week tooth germ had significantly larger proportions of follicle cells, and significantly smaller proportion of distal pulps (Figure 4C).

**Figure 4.**
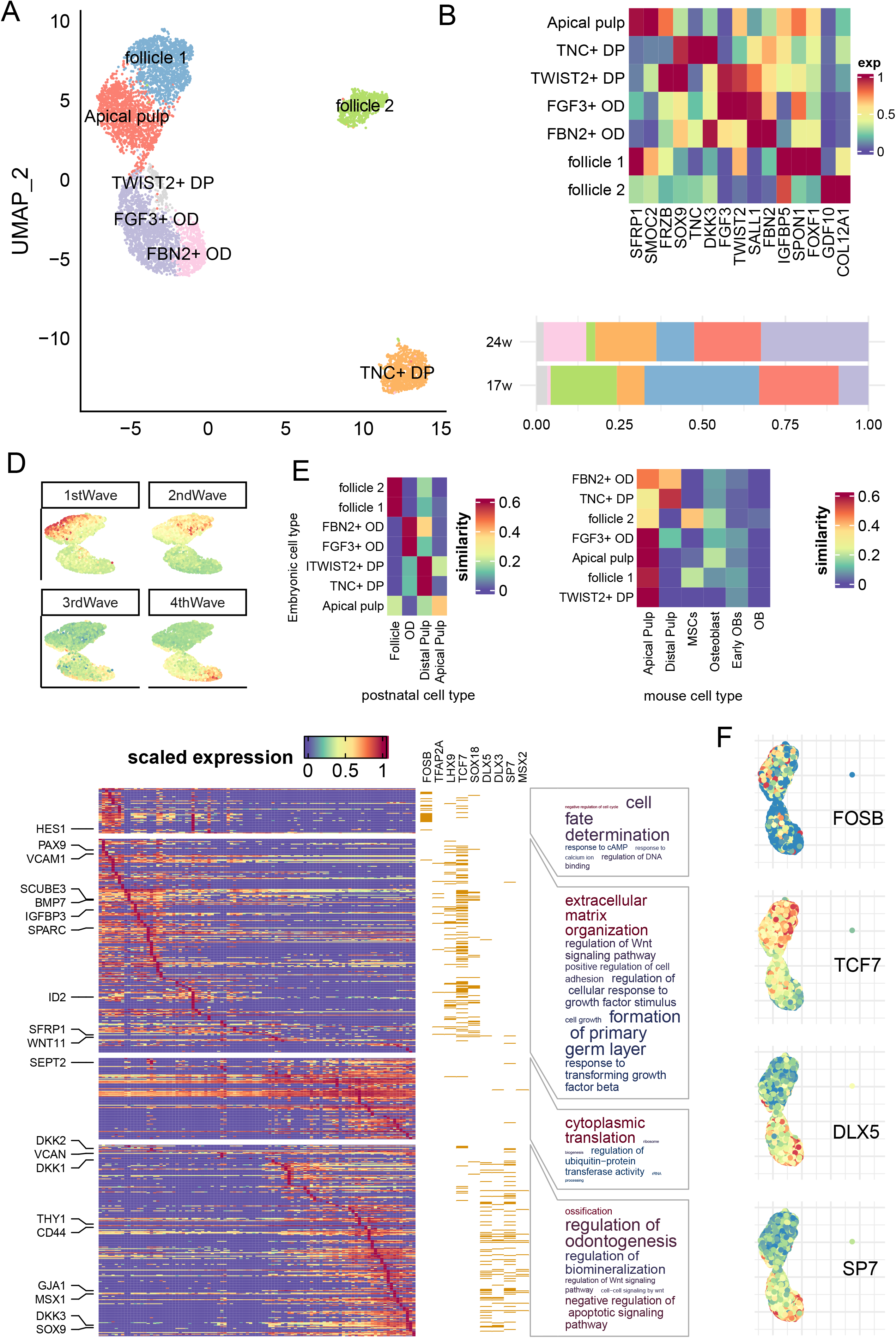
In-depth analysis of dental mesenchyme. A -C: similar to Figure 2 A-C, but for epithelium clusters. Pseudotime trajectory was shown in Figure S7. OD: pre-odontoblast. AP: apical progenitor. MSC: mesenchymal stem cell. D: Expression of four waves of gene along mesenchyme development. E: Heatmap of module gene expression along pseudotime. Yellow bar denoted that the gene was predicted to be target of the corresponding transcription factor. Word cloud showed GO term enriched by each wave. Font size denoted enrichment odds ratio, color denoted enrichment p value. F: Expression of selected regulons.

By integrating our data with previous dental cell atlas from Hermans et al.[13] (Figure 4E), we found that human embryonic dental mesenchyme was generally similar to postnatal cells: follicle\ odontoblast and distal pulp subgroups were all closest to the same cluster in postnatal data. The only exception was apical pulp, which was moderately distinct from postnatal apical pulp. Nonetheless, when comparing human embryo to mouse embryo, we found substantial difference. In fact, only TNC+ DP resembled the corresponding DP cluster in mouse data, and all other cell types were close to mouse apical pulp.

We further applied pseudotime analysis to reconstruct the developmental trajectory on the continuous cell clusters from follicle 1 to FBN2+ OD (Figure S7), and identified four waves of gene expression patterns along this trajectory (Table S8). As shown in Figure 4D, the first and second wave corresponded to progenitor cell differentiation, and the third and fourth wave corresponded to pulp cell development. The first wave included HES1 and TWIST1, and were enriched in cell fate determination (GO p-adjust=2.28×10^−3^) and other processes related to cell cycle (Figure 4E). SCENIC[18] analysis revealed that TFs regulating the first wave genes were immediate genes like FOS and JUN family like FOSB (Fisher test p=3.60 × 10^−14^; Figure 4E and Table S9). The second wave included genes like PAX9, SCUBE3 and SPARC, and were enriched in extracellular matrix organization (GO p-adjust=1.23×10^−18^) and formation of primary germ layer (GO p-adjust=8.38×10^−9^). They were also enriched in the predicted targets of LHX9, TCF7 and SOX18 (Fisher test p<10^−10^; Figure 4F and Table S9). The third wave consisted of genes taking part in cytoplasmic translation (GO p-adjust=7.66×10^−29^), without significant enrichment in specific TF regulon. Lastly, the fourth wave genes, including DKK2, VCAN and THY1, were involved in regulation of odontogenesis (GO p-adjust=1.68 × 10^−4^), regulation of Wnt signaling pathway (GO p-adjust=0.0001) and ossification (GO p-adjust=3.18×10^−6^). They were mainly regulated by TF like DLX5,DLX3, SP7 and MSX2 (Fisher test p<1.78×10^−5^; Figure 4F and Table S9). The overall expression levels of regulons of these highlighted TF were also in line with the four wave patterns (Figure 4F).

### Pulp cell subtypes had distinct spatial distribution

We applied Seurat anchor-based integration algorithm[15] to map single cell data to spatial transcriptome (Figure 5A). In 17-week primary tooth germ, we observed a follicle→distal pulp→apical pulp→epithelium layer distribution from distal to proximal end, in line with their order in the developmental trajectory (Figure 4A). In the 24-week primary tooth germ, we further discovered that FBN2+ OD was at the outer layer of pulp cell, which consisted of TWIST2+ DP and FGF3+ OD. One exception was the TNC+ DP, which was sparsely distributed in permanent tooth germ without significant spatial structure (Figure 5A and Figure S8). We plotted the expression of its marker TNC and HEY1 and also found no matching patterns (Figure 5B). We inferred that TNC+ DP was loosely distributed in all regions of pulp tissues. As a comparison, the density as well as marker expression of other mesenchyme cell subtypes (Figure 5B and Figure S8 & Figure S9) were all aggregated to specific regions. The temporal distribution (Figure 5C) showed that TNC+ DP and two follicle and subgroups were mainly found in immature permanent tooth germ (p<10^−10^), whereas odontoblast and pulp cells were mainly found in mature primary tooth germ (p<10^−10^).

**Figure 5.**
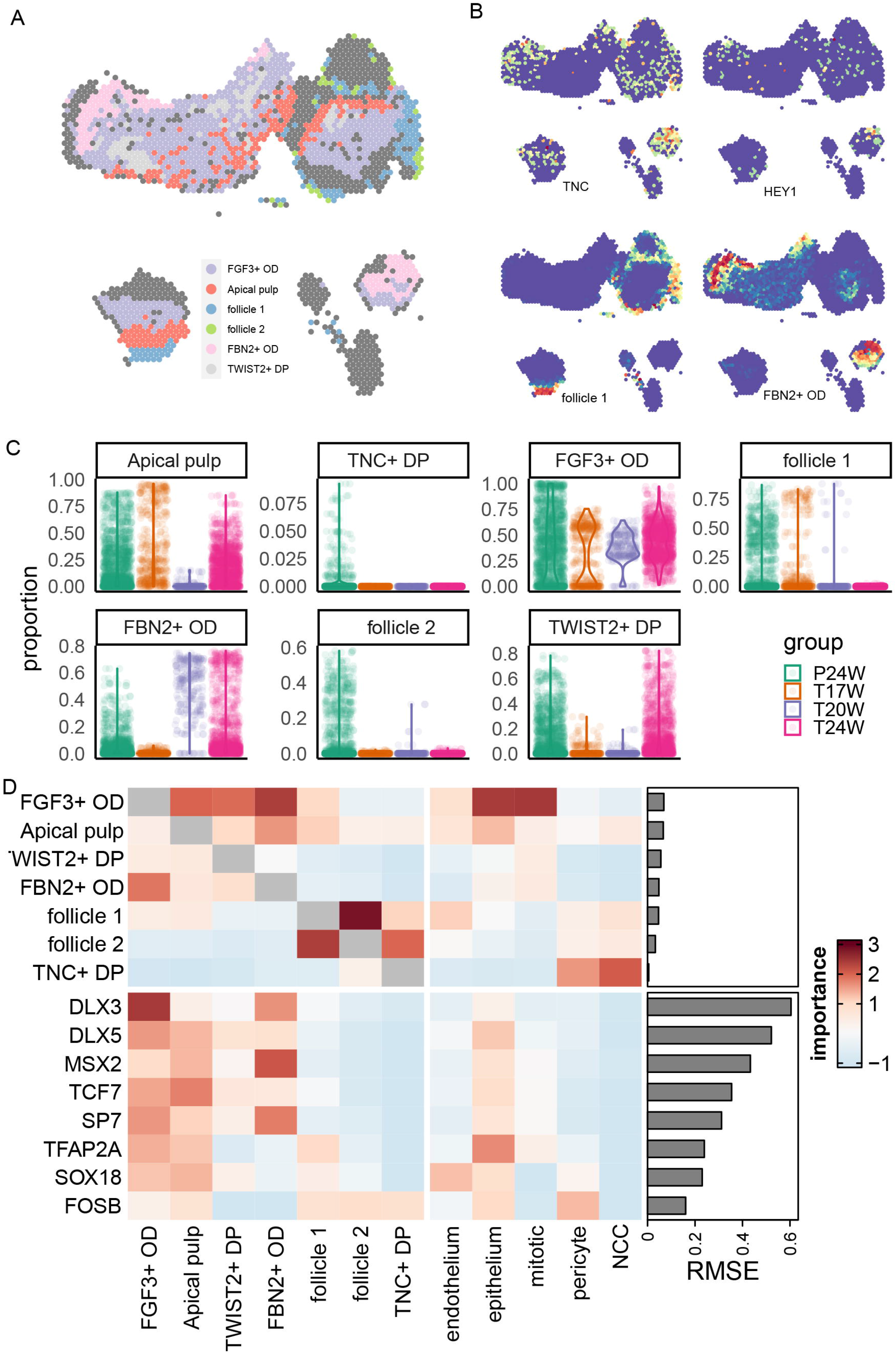
Spatial transcriptome of dental mesenchyme. A: Cell type annotation of each spot. B: Cell type probability spatial distribution and marker gene spatial expression. C: Cell type probability distribution per developmental stage. D: misty result of spatial adjacency, quantified by model importance score. RMSE: Root mean squared error of each model.

We also applied misty[21] to quantify the spatial adjacency among cell subtypes as well as the key TFs. Within mesenchyme cell groups, we found close relationship between two subgroups of follicle cells (misty adjacency score>2.5) as well as FGF3+ OD and FBN2+ OD (adjacency score>2). Outside mesenchyme cell groups, only epithelium and mitotic cells had close relationship with pulp cells (adjacency score>2). For TFs, pulp cells had close association with nearby expression values of multiple TFs that had important roles in mesenchyme differentiation, except FOSB which took effect in early progenitors. Another spatial association was found between FBN2+ OD and MSX2 (adjacency score=2.02).

### Signaling network analysis of dental development

Having discovered the refined cell subtypes of embryonic tooth germ, we were able to study the intercellular signaling pathways that regulate dental development. We first applied nichenetr[20] to build a signaling network that regulate branch-specific key genes in epithelial development (181 key genes included in nichenetr; Figure S10). Integrating the prioritized upstream ligands of both root branch, SI branch and ameloblast branch, we found that signals from mesenchyme (including follicle cell) and pericyte regulated the largest number of epithelial key genes (n=142 and 92, respectively, Figure S10 and Table S10). Top credible ligands included BMP5, SFRP2, JAG1, HMGB2 (Table S11), etc. Among them, JAG1 was predicted to regulate the largest number (n=91) of epithelial key genes, including GJA1, NOTCH1, etc. Two notable exceptions were GPNMB and SHH: they were the only prioritized ligands that showed highest expression in ameloblast (Figure S11) and had high potential to regulate SI and ameloblast branch development (AUC=0.59 and 0.53, respectively), suggesting that they might act through autocrine or paracrine. Despite their expression in epithelium, the highest expression was found in SCP (Figure S10).

Lastly, we analyzed the intercellular signaling networks that regulated dental mesenchyme development. For all 162 mesenchyme key genes (genes of first, second and fourth wave of Figure 4D), 135 were regulated by ligands from pericyte, and 133 were regulated by ligands from epithelium (Figure S12). These included ligands like BMP5, BMP7, DSC3 and COL4A1, etc. We also found that different ligands tended to regulate different stages of mesenchyme development. For example, BMP5 and BMP7 regulated all first wave genes and more than 70% of the second wave genes that were included in nichenetr analysis, whereas all predicted targets of APP (mainly expressed in endothelium) and TGFB1 (mainly expressed in macrophage) belonged to the fourth wave (Figure S12). In general, upstream signaling pathways of mesenchyme development was restricted to a few vital ligands and cell types, such as BMP signals from pericyte and epithelium.

We also applied Misty to quantify whether these signaling network were spatially adjacent to their targets, as detailed in Supplementary materials.

## Discussion

In the current study, we applied single-cell RNA sequencing and spatial transcriptome on human embryonic tooth germ to depict a spatiotemporal cell landscape of human tooth development. We revealed the cellular composition as well as their spatial distribution in human embryonic tooth germ spanning different developmental periods and tissues from both primary and permanent tooth germ. We also depicted the developmental trajectory, biological insights, essential transcription factors and signaling pathways of dental epithelium and mesenchyme development. In complement with the existing dental cell atlas of prenatal and postnatal mouse tooth[6, 7], postnatal human tooth[7, 9] and tooth germ[12], our result brought one more step closer to the intact single-cell picture of dental development[13].

Our collection of human embryonic dental tissues has filled in the blank of cell atlas of early development process of human tooth. Currently available single-cell data on human tooth[13] came from early youth or child volunteers, which inevitably missed progenitor cells at early stage. On the other hand, studies based on mouse embryonic tissues might not be immediately translatable to human research due to profound cross-species difference. Rodent incisor has a relatively short lifespan and routinely lost. Many cell types, including ameloblasts and odontoblasts that secrete mineralized enamel and dentin, respectively, are continuously proliferated from by self-renewal cells located in the proximal part of the tooth[23]. Some of the vital structures of this process, such as the transit-amplifying (T-A) zone, were not found in human dental tissue[24]. Thus, it is desirable to generate the dental cell landscape for both human and mice at the developing stage to achieve intact understanding of human dental generation.

Our result revealed that epithelial cells and mesenchyme cells exhibited profound crosstalk and regulated the developmental process of each other via intercellular signaling networks[25]. In mouse incisor, BMP-SMAD-SHH signals from a transient structure Hartwig’s epithelial root sheath[26] interacts with mesenchymal cells and regulated root elongation and epithelium maturation[27]. Epithelium could also regulate the mineralization of nearby dentin though secretion of signal molecules[28], including Wnt and fibroblast growth factors[29]. In line with these observations, previous scRNA intercellular signaling network analysis on postnatal pulp[10] and tooth germ[12] tissues have demonstrated the existence of dense communication. Our work extended these knowledges to prenatal epithelium-mesenchyme developmental process and highlighted more signal pathways like JAG1, SFRP2 and COL4A1.

In this study, we focus on a narrow development window (17-24 PCW). Enamel organs mature during the 12th week of embryonic development, and then the odontoblasts start forming the dentin. We selected fetal tooth germs from weeks 17 to 24 because tooth germs of younger gestational age are not yet fully developed and are relatively small in size, making it difficult to obtain samples. On the other hand, tooth germs of older gestational age already have a considerable amount of dental hard tissues formed, and during sampling, it is necessary to remove these hard tissues, resulting in the loss of a portion of the tooth germ tissue.

Our study has limitations. Earlier (<15 gestational weeks) and later (>30 gestational weeks) embryonic healthy tissues are difficult to access due to ethnic concerns and other issues discussed above, and dental developmental process at these stages could not be covered in the current study. The spatial structures at the third dimension are also not complete in different slicing procedures. Future efforts are desired to fill in these blanks.

In conclusion, we constructed a spatiotemporal cell landscape of human embryonic tooth germ, depicted developmental trajectory of dental epithelium and mesenchyme, and revealed the biological mechanisms underlying these processes. Our result served as a foundation for future research on dental development and stem cell engineering.

## Acknowledgement

This work was supported by Research Fund of Shanghai Tongren Hospital, Shanghai Jiaotong University School of Medicine (No: TRYJ2021JC14).

## Author contribution

Y.S designed the study, W.W and J.Q supervised the study. J.Q and L.H collected and preprocessed the sample. Y.S, Y.Y, S.W, K.G, J.Y, J.L and S.S performed the experiments. Y.S analyzed the data and drafted the manuscript. All authors read, revised and approved the manuscript.

## Data availability statement

Data could be downloaded at 10.17632/v3wgx8pm5y.1.

## Reference

1. Yu T, Klein OD (2020) Molecular and cellular mechanisms of tooth development, homeostasis and repair. Dev 147:. https://doi.org/10.1242/DEV.184754/225112

2. Balic A (2019) Concise Review: Cellular and Molecular Mechanisms Regulation of Tooth Initiation. Stem Cells 37:26–32

3. Thesleff I (2014) Current understanding of the process of tooth formation: Transfer from the laboratory to the clinic. Aust Dent J 59:48–54. https://doi.org/10.1111/adj.12102

4. Matalová E, Lungová V, Sharpe P (2015) Development of Tooth and Associated Structures. In: Stem Cell Biology and Tissue Engineering in Dental Sciences. Elsevier Inc., pp 335–346

5. Wu J, Ding Y, Wang J, et al (2022) Single-cell RNA sequencing in oral science: Current awareness and perspectives. Cell Prolif 55:e13287. https://doi.org/10.1111/CPR.13287

6. Jing J, Feng J, Yuan Y, et al (2022) Spatiotemporal single-cell regulatory atlas reveals neural crest lineage diversification and cellular function during tooth morphogenesis. Nat Commun 13:1–14. https://doi.org/10.1038/s41467-022-32490-y

7. Krivanek J, Soldatov RA, Kastriti ME, et al (2020) Dental cell type atlas reveals stem and differentiated cell types in mouse and human teeth. Nat Commun 11:1–18. https://doi.org/10.1038/s41467-020-18512-7

8. Chiba Y, Saito K, Martin D, et al (2020) Single-Cell RNA-Sequencing From Mouse Incisor Reveals Dental Epithelial Cell-Type Specific Genes. Front Cell Dev Biol 8:841. https://doi.org/10.3389/fcell.2020.00841

9. Pagella P, de Vargas Roditi L, Stadlinger B, et al (2021) A single-cell atlas of human teeth. iScience 24:102405. https://doi.org/10.1016/J.ISCI.2021.102405

10. Yin W, Liu G, Li J, Bian Z (2021) Landscape of Cell Communication in Human Dental Pulp. Small methods 5:. https://doi.org/10.1002/SMTD.202100747

11. Lee S, Chen D, Park M, et al (2022) Single-Cell RNA Sequencing Analysis of Human Dental Pulp Stem Cell and Human Periodontal Ligament Stem Cell. J Endod 48:240–248. https://doi.org/10.1016/J.JOEN.2021.11.005

12. Shi Y, Yu Y, Zhou Y, et al (2021) A single-cell interactome of human tooth germ from growing third molar elucidates signaling networks regulating dental development. Cell Biosci 11:1–17. https://doi.org/10.1186/S13578-021-00691-5

13. Hermans F, Bueds C, Hemeryck L, et al (2022) Establishment of inclusive single-cell transcriptome atlases from mouse and human tooth as powerful resource for dental research. Front Cell Dev Biol 10:1990. https://doi.org/10.3389/FCELL.2022.1021459/BIBTEX

14. Wang B, Li H, Liu Y, et al (2014) Expression patterns of WNT/β-CATENIN signaling molecules during human tooth development. J Mol Histol 45:487–496. https://doi.org/10.1007/S10735-014-9572-5/FIGURES/5

15. Stuart T, Butler A, Hoffman P, et al (2019) Comprehensive Integration of Single-Cell Data. Cell 177:1888–1902.e21. https://doi.org/10.1016/j.cell.2019.05.031

16. Tirosh I, Izar B, Prakadan SM, et al (2016) Dissecting the multicellular ecosystem of metastatic melanoma by single-cell RNA-seq. Science (80-) 352:189–196. https://doi.org/10.1126/SCIENCE.AAD0501/SUPPL_FILE/TIROSH.SM.PDF

17. Cao J, Spielmann M, Qiu X, et al (2019) The single-cell transcriptional landscape of mammalian organogenesis. Nature 566:496–502. https://doi.org/10.1038/s41586-019-0969-x

18. Aibar S, González-Blas CB, Moerman T, et al (2017) SCENIC: single-cell regulatory network inference and clustering. Nat Methods 14:1083–1086. https://doi.org/10.1038/nmeth.4463

19. Yu G, Wang L-G, Han Y, He Q-Y (2012) clusterProfiler: an R Package for Comparing Biological Themes Among Gene Clusters. Omi A J Integr Biol 16:284–287. https://doi.org/10.1089/omi.2011.0118

20. Browaeys R, Saelens W, Saeys Y (2020) NicheNet: modeling intercellular communication by linking ligands to target genes. Nat Methods 17:159–162. https://doi.org/10.1038/s41592-019-0667-5

21. Tanevski J, Flores ROR, Gabor A, et al (2022) Explainable multiview framework for dissecting spatial relationships from highly multiplexed data. Genome Biol 23:1–31. https://doi.org/10.1186/S13059-022-02663-5/FIGURES/7

22. Sharir A, Marangoni P, Zilionis R, et al (2019) A large pool of actively cycling progenitors orchestrates self-renewal and injury repair of an ectodermal appendage. Nat Cell Biol 21:1102–1112. https://doi.org/10.1038/s41556-019-0378-2

23. Seidel K, Marangoni P, Tang C, et al (2017) Resolving stem and progenitor cells in the adult mouse incisor through gene coexpression analysis. Elife 6:. https://doi.org/10.7554/ELIFE.24712

24. Kuang-Hsien Hu J, Mushegyan V, Klein OD (2014) On the cutting edge of organ renewal: Identification, regulation, and evolution of incisor stem cells. genesis 52:79–92. https://doi.org/10.1002/DVG.22732

25. Puthiyaveetil JSV, Kota K, Chakkarayan R, et al (2016) Epithelial – Mesenchymal Interactions in Tooth Development and the Significant Role of Growth Factors and Genes with Emphasis on Mesenchyme – A Review. J Clin Diagn Res 10:ZE05. https://doi.org/10.7860/JCDR/2016/21719.8502

26. Huang X, Xu X, Bringas P, et al (2010) Smad4-Shh-Nfic signaling cascade-mediated epithelial-mesenchymal interaction is crucial in regulating tooth root development. J Bone Miner Res 25:1167–1178. https://doi.org/10.1359/JBMR.091103

27. Li J, Feng J, Liu Y, et al (2015) BMP-SHH Signaling Network Controls Epithelial Stem Cell Fate via Regulation of Its Niche in the Developing Tooth. Dev Cell 33:125–135. https://doi.org/10.1016/j.devcel.2015.02.021

28. Lavicky J, Kolouskova M, Prochazka D, et al (2022) The Development of Dentin Microstructure Is Controlled by the Type of Adjacent Epithelium. J Bone Miner Res 37:323–339. https://doi.org/10.1002/JBMR.4471

29. Hermans F, Hemeryck L, Lambrichts I, et al (2021) Intertwined Signaling Pathways Governing Tooth Development: A Give-and-Take Between Canonical Wnt and Shh. Front cell Dev Biol 9:. https://doi.org/10.3389/FCELL.2021.758203

