## Supplementary for "Spatiotemporal cell landscape of human embryonic tooth development": supplementary.docx (from another version of this preprint: version 1, 2023-03-02)

### Supplementary Methods

Samples used for scRNA were minced on ice to less than 1 mm cubic pieces, followed by enzymatic digestion using Sigma with manual shaking every 5 min. Samples were then centrifuged at 300 rcf for 30 sec at room temperature and removed the supernatant without disturbing the cell pellet. Next, 1× PBS (calcium and magnesium free) containing 0.04% weight/volume BSA (400 µg/ml) was added and then centrifugation at 300 rcf for 5 min. The cell pellet was resuspended in 1 ml red blood cell lysis buffer and incubated for 10 min at 4℃. After Red Blood Cell Lysis, samples were resuspended in 1 ml PBS containing 0.04% BSA. Next, samples were filtered over Scienceware Flowmi 40-µm cell strainers (VWR). After tumor dissociation, cell concentration and cell viability were determined by hemocytometer and Trypan Blue staining.

Cellular suspensions were loaded on a Chromium Single Cell Controller instrument (10x Genomics, CA, USA) to generate single-cell GEMs. Single-cell RNA-Seq libraries were prepared using the Chromium Single Cell 5’ Library & Gel Bead Kit (P/N 1000006, 10x Genomics). GEM-RT was performed in a C1000 Touch Thermal cycler with 96-Deep Well Reaction Module (Bio- Rad; P/N 1851197): 53℃ for 45 min, 85℃ for 5 min; held at 4℃, then broken and the single-strand cDNA was cleaned up with DynaBeads MyOne Silane Beads (Thermo Fisher Scientific; P/N 37002D). Barcoded, full length cDNA was amplified using the C1000 Touch Thermal cycler with 96-Deep Well Reaction Module: 98℃ for 45 s; cycled 13 × : 98℃ for 20 s, 67℃ for 30 s, and 72℃ for 1 min; 72 ℃ for 1 min; held at 4℃. Amplified cDNA product was cleaned up with the SPRIselect Reagent Kit (0.6 × SPRI; Beckman Coulter; P/N B23318). Barcoded, full-length V(D)J segments were enriched from amplified cDNA with primers specific to TCR constant regions. The target enrichment 1 was performed in a C1000 Touch Thermal cycler with 96-Deep Well Reaction Module: 98℃ for 45 s; cycled 10 × : 98℃ for 20 s, 67℃ for 30 s, and 72℃ for 1 min; 72℃ for 1 min; held at 4℃. The target enrichment 1 was cleaned up with the SPRIselect Reagent Kit (0.8 × SPRI). The target enrichment 2 was performed in a C1000 Touch Thermal cycler with 96-Deep Well Reaction Module: 98℃ for 45 s; cycled 10 × : 98℃ for 20 s, 67℃ for 30 s, and 72℃ for 1 min; 72℃ for 1 min; held at 4℃. The target enrichment 2 was cleaned up twice with the SPRIselect Reagent Kit (0.5 × and 0.8 × SPRI). 5’ gene expression and enriched libraries were constructed using the reagents in the Chromium Single Cell 3’/5’ Library Construction kit (P/N 1000020). For 5’ gene expression library construction, these steps were followed: (1) fragmentation, end repair and A-tailing; (2) post fragmentation, end repair and A-tailing cleanup with SPRIselect; (3) adaptor ligation; (4) post ligation cleanup with SPRIselect; (5) sample index PCR and cleanup. For the enriched library construction, these steps were followed: (1) fragmentation, end repair and A-tailing; (2) adaptor ligation; (3) post ligation cleanup with SPRIselect; (4) sample index PCR and cleanup.

Tooth germ samples were cryosectioned coronally to a thickness of 10 μm (bregma: − 2.52 to − 2.92, interaural: 6.08 to 6.48 mm) using a CryoStar NX70 cryostat (Thermo Fisher Scientific, Waltham, MA, USA). Tissue sections were layered onto the visium spatial tissue optimization slide containing oligonucleotides for mRNA capture (10× Genomics, Pleasanton, CA, USA). Each capture area per slide has 5000 spatially barcoded with a diameter of 55 μm and a center-to-center distance of 100 μm, over an area of 6.5 mm by 6.5 mm. A Master Mix containing reverse transcription (RT) reagents and fluorescently labeled nucleotides is added on top of the tissue sections, resulting in fluorescently labeled cDNA synthesis. Tissue is enzymatically removed, leaving behind fluorescent cDNA covalently linked to oligonucleotides on the slide. Fluorescent cDNA is visualized under fluorescence imaging conditions verified using the Visium Imaging Test Slide. Fluorescent cDNA is visualized under fluorescence imaging conditions verified using the Visium Imaging Test Slide. H&E and fluorescence images are compared. The permeabilization time that results in maximum fluorescence signal with the lowest signal diffusion is optimal. The libraries were sequenced with paired-end 150 bp sequencing (PE150) by NovaSeq 6000 platform.

Space ranger showed the capture area of the tissue in the slide and differentiates reads for each spot based on spatial barcode information. STAR was used to assess the sample quality by total number of spots, the number of pairs of reads in each Spot, the number of detected genes, and the number of unique molecular identifier (UMIs).

The Cell Ranger software pipeline (version 5.0.0) provided by 10×Genomics was used to demultiplex cellular barcodes, map reads to the genome and transcriptome using the STAR aligner, and down-sample reads as required to generate normalized aggregate data across samples, producing a matrix of gene counts versus cells. We processed the unique molecular identifier (UMI) count matrix using the R package Seurat (version 3.1.1). To remove low quality cells and likely multiplet captures, which is a major concern in microdroplet-based experiments, we applied a criteria to filter out cells with UMI/gene numbers out of the limit of mean value +/- 2 fold of standard deviations assuming a Guassian distribution of each cells' UMI/gene numbers. Following visual inspection of the distribution of cells by the fraction of mitochondrial genes expressed, we further discarded low-quality cells where >10% of the counts belonged to mitochondrial genes. Additionally, we applied DoubletFinder package[2] (version 2.0.2) to identify potential doublet. Top variable genes across single cells were identified using the method described in Macosko et al[3]. The most variable genes were selected using FindVariableGenes function(mean.function = FastExpMean, dispersion.function = FastLogVMR) in Seurat[1]. Principal component analysis (PCA) was performed to reduce the dimensionality with RunPCA function in Seurat[1].

### Supplementary Figure S1

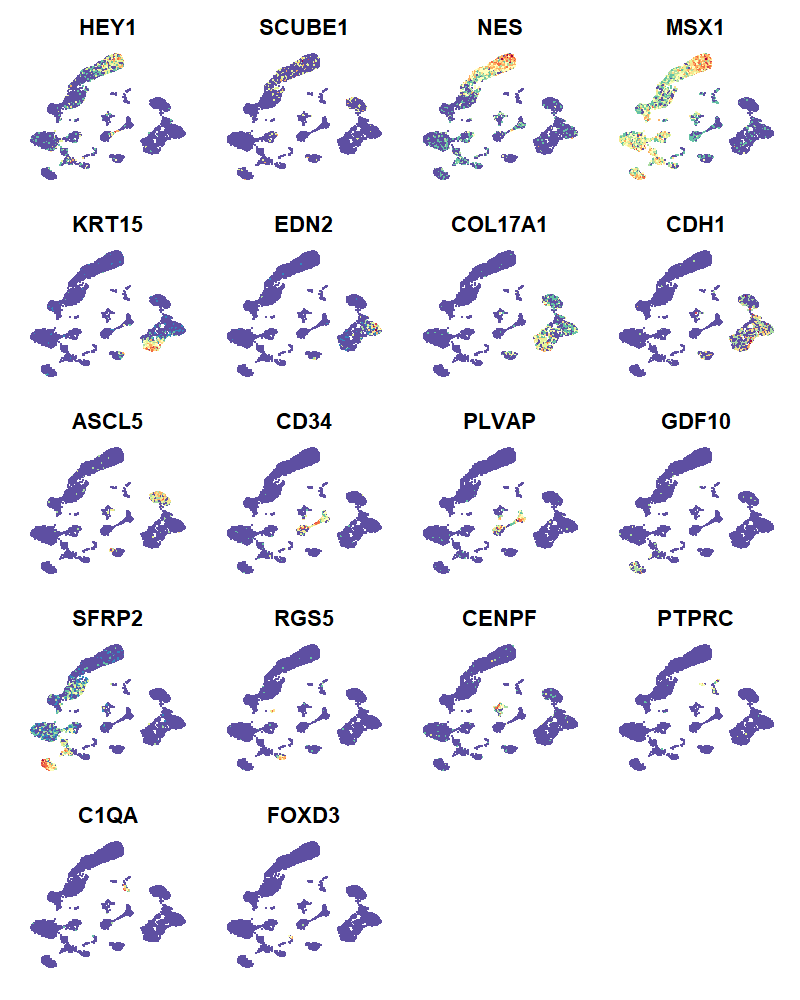

Marker expression of major cell types in single cell data.

### Supplementary Figure S2

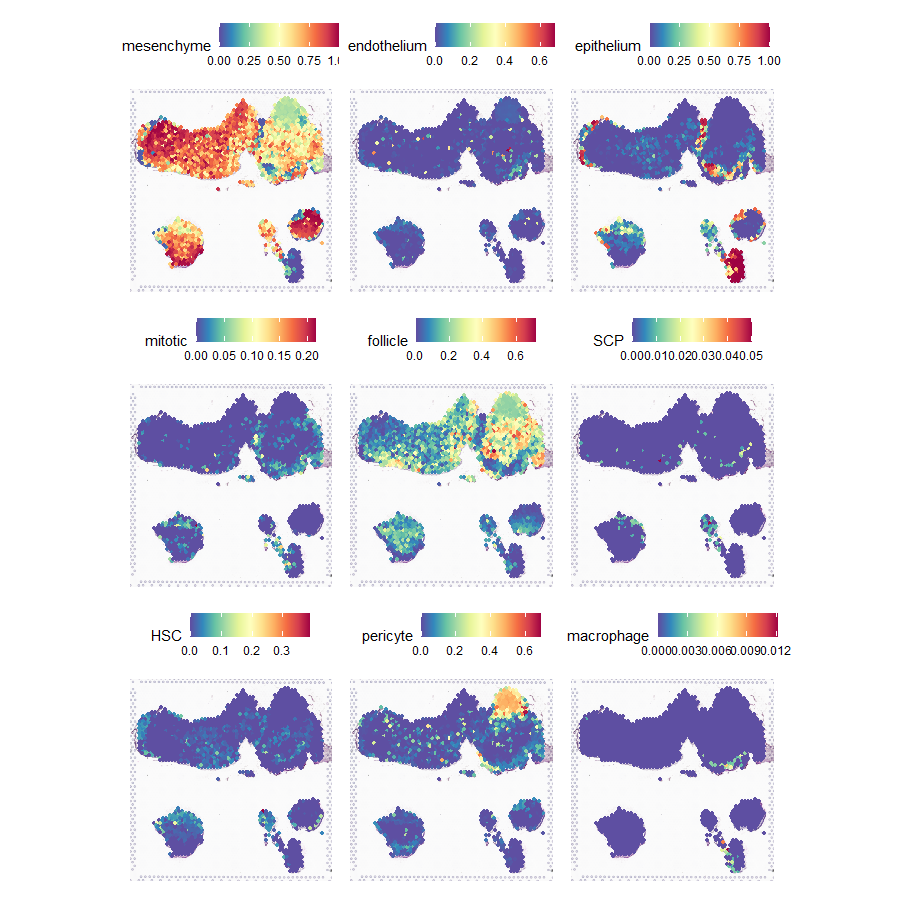

Marker expression of major cell types in Spatial transcriptome data.

### Supplementary Figure S3

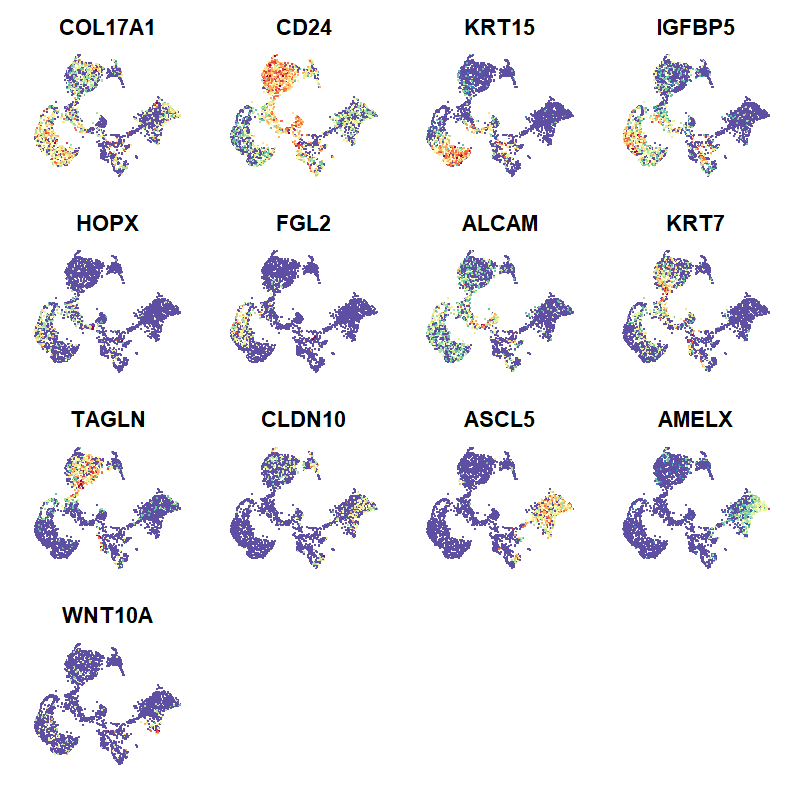

Marker expression of epithelial subtypes in single cell data.

### Supplementary Figure S4

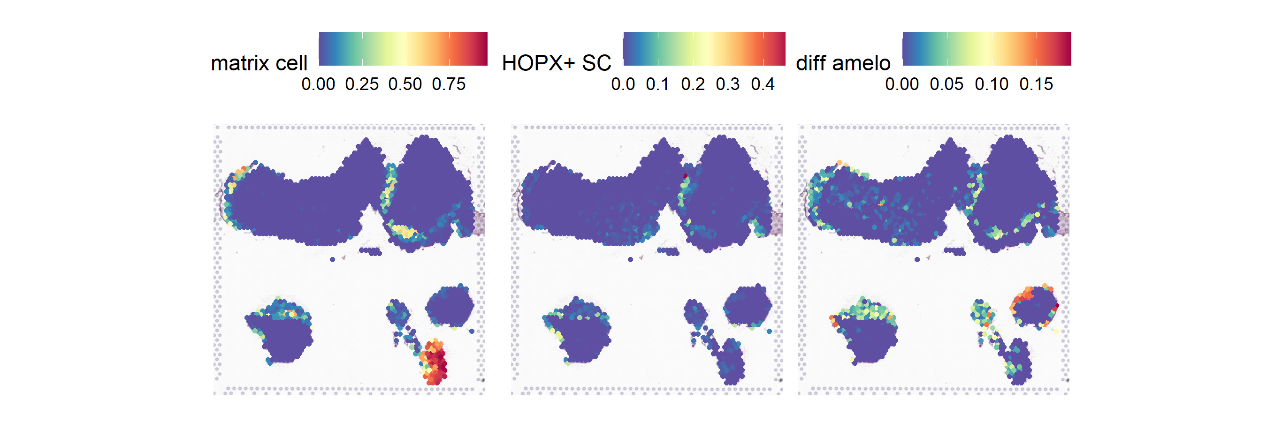

Cell type proportion of epithelial subtypes in spatial transcriptome data.

### Supplementary Figure S5

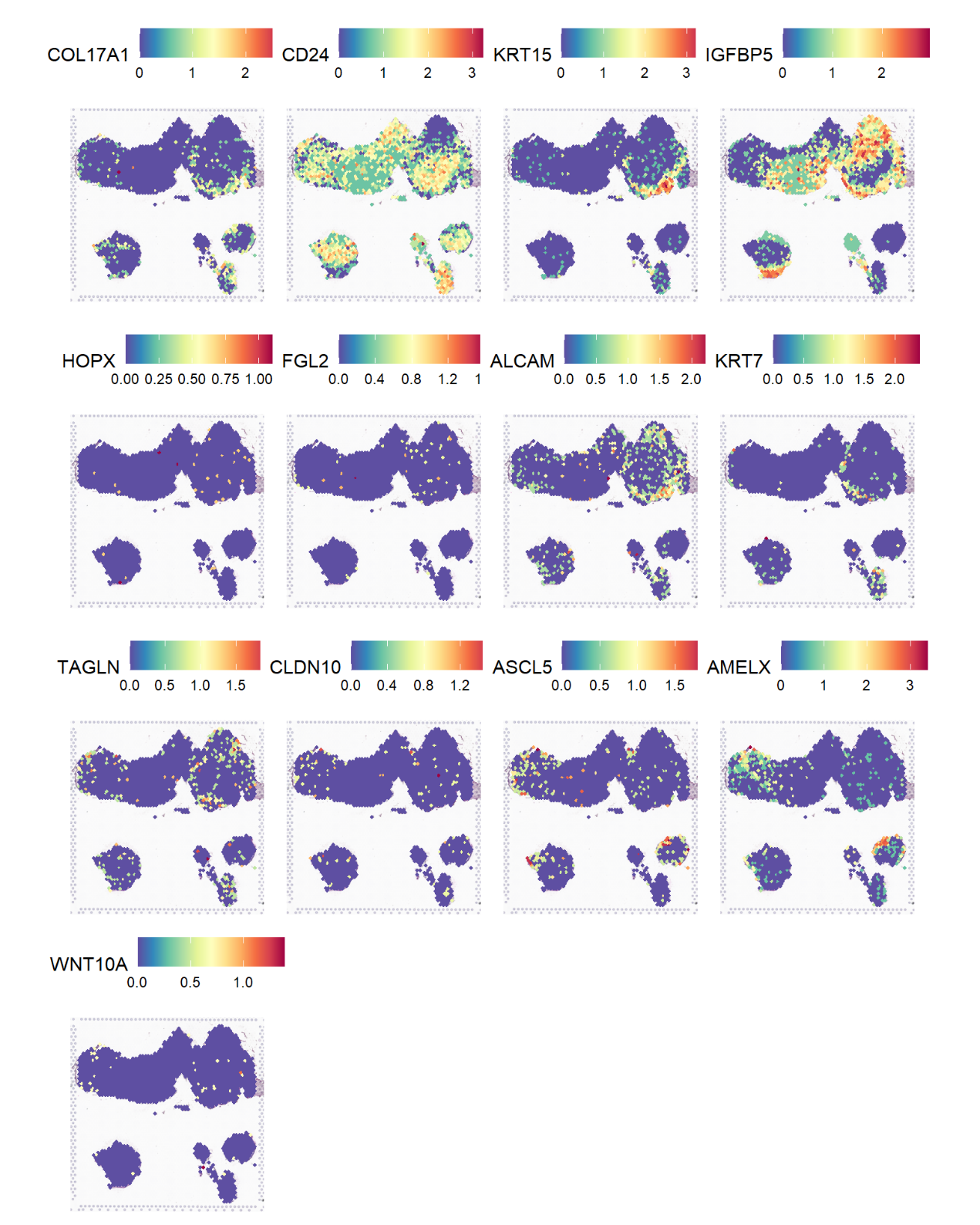

Marker expression of epithelial subtypes in spatial transcriptome data.

### Supplementary Figure S6

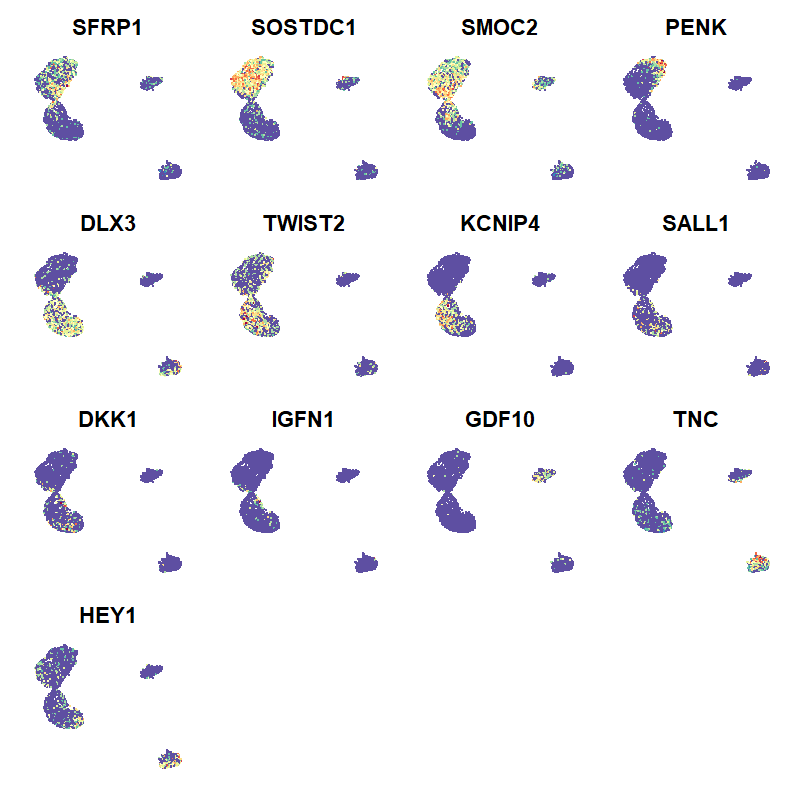

Marker expression of mesenchyme subtypes in single cell data.

### Supplementary Figure S7

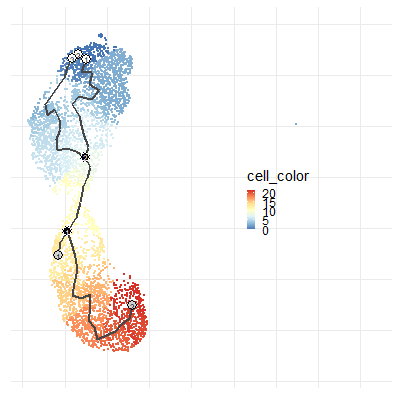

Pseudotime trajectory of mesenchyme development.

### Supplementary Figure S8

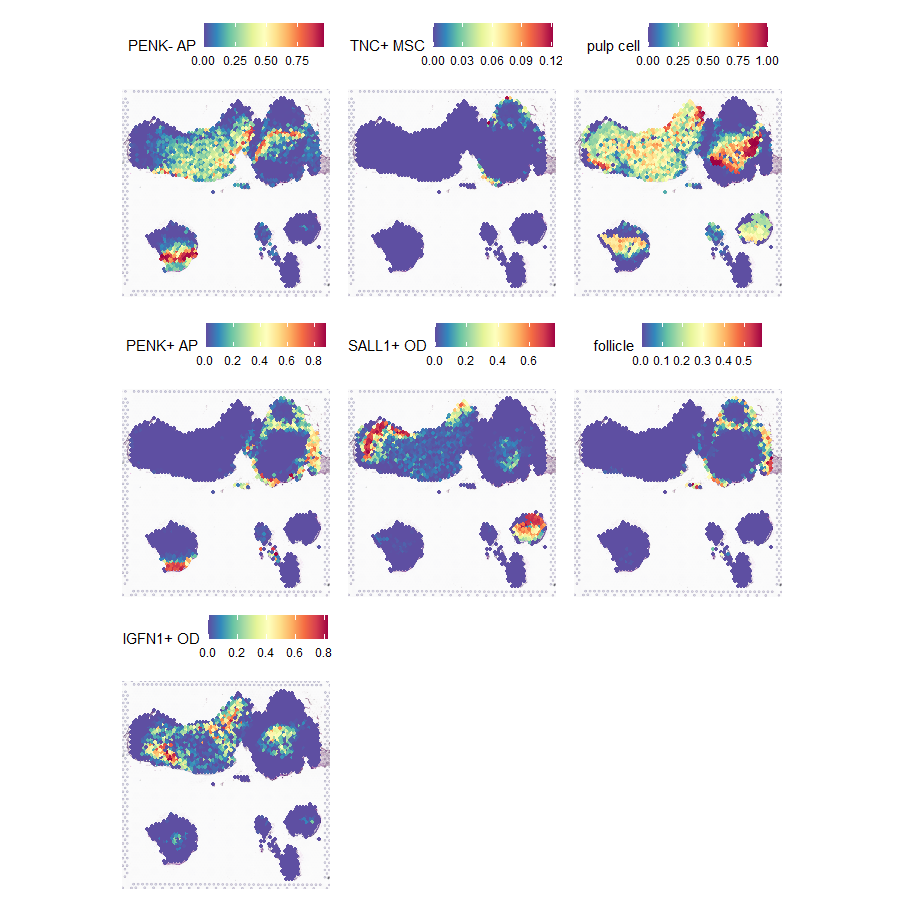

Cell proportion of mesenchyme subtypes in spatial transcriptome data.

### Supplementary Figure S9

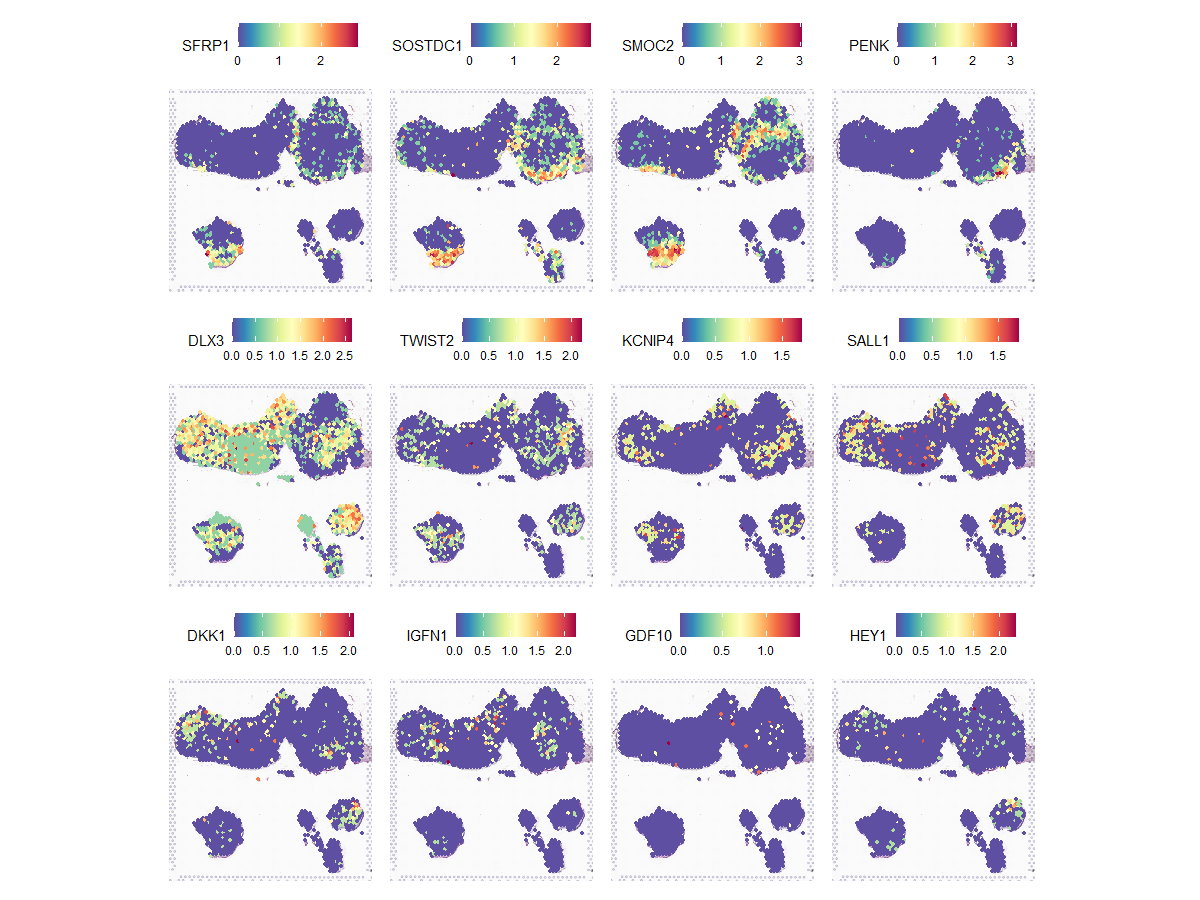

Marker expression of mesenchyme subtypes in spatial transcriptome data.

### Supplementary Figure S10

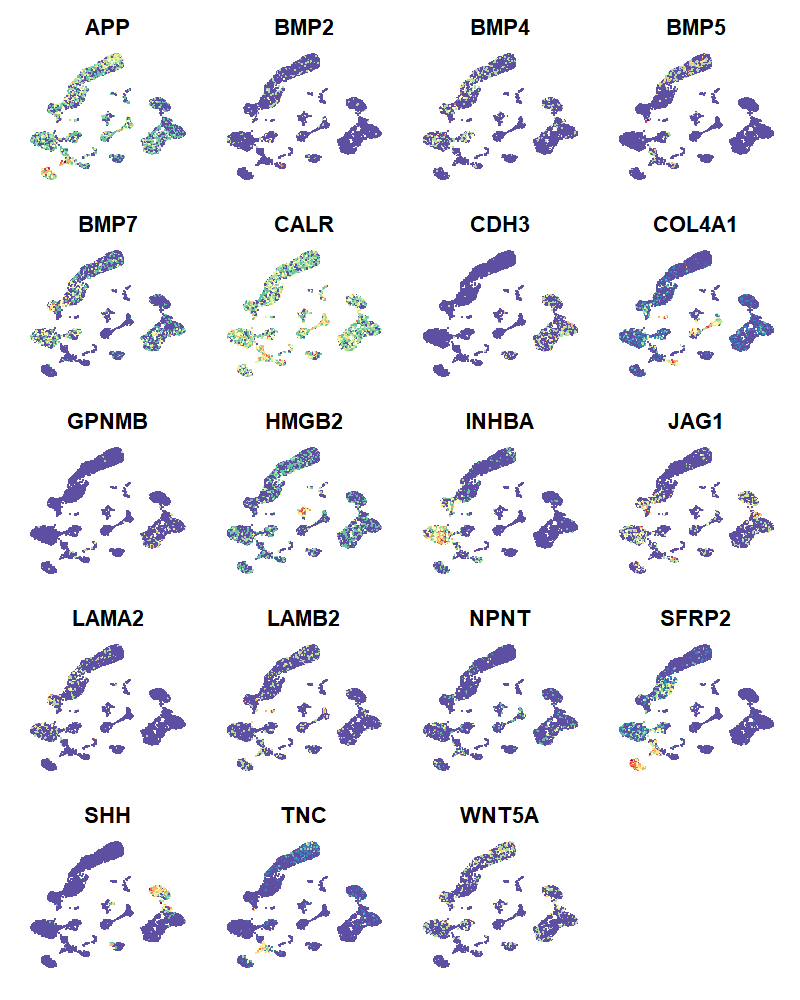

Expression of selected signaling molecules of epithelial development in single cell data.

### Supplementary Figure S11

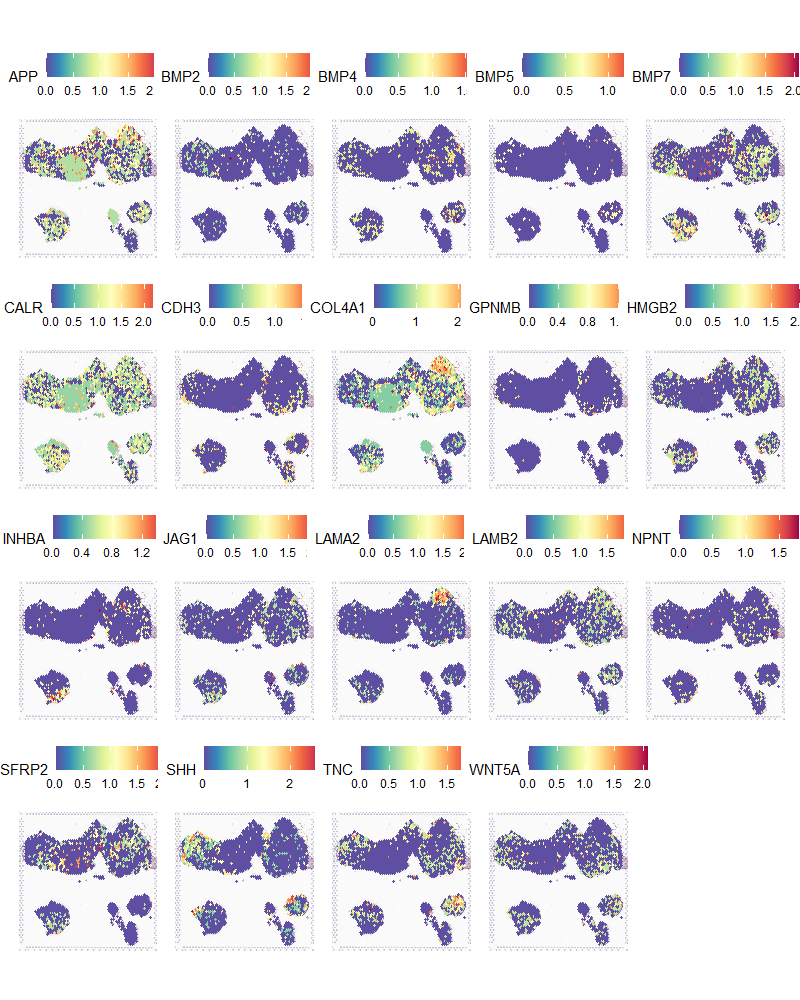

Expression of selected signaling molecules of epithelial development in spatial transcriptome data.

### Supplementary Figure S12

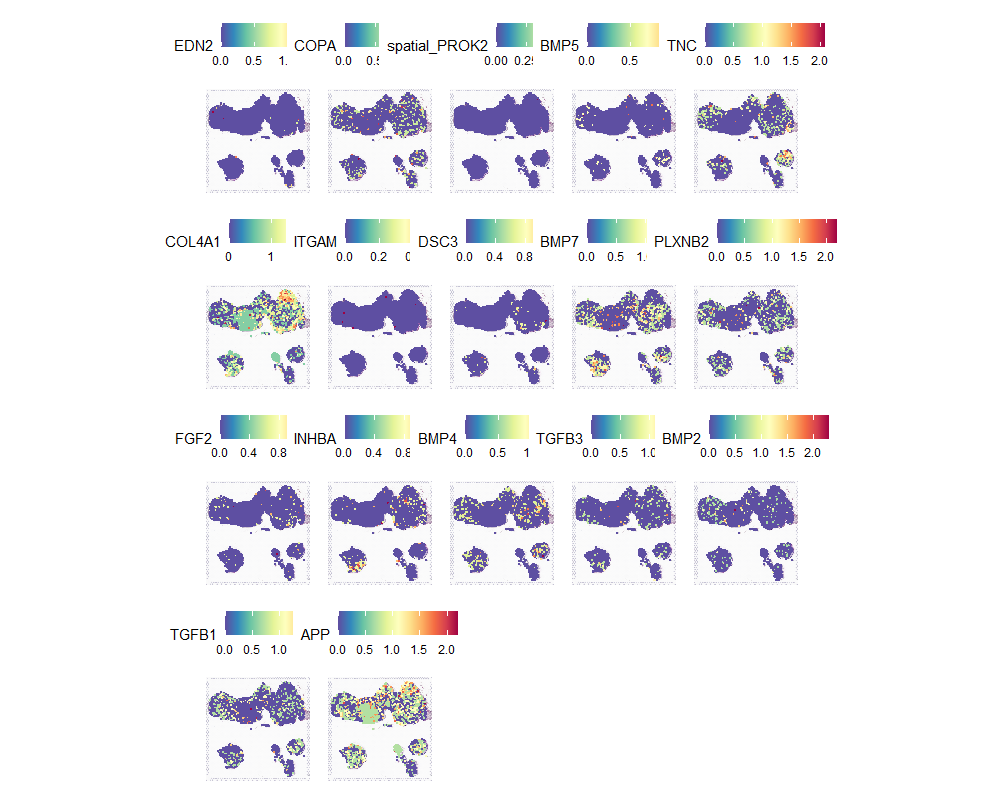

Expression of selected signaling molecules of mesenchyme development in spatial transcriptome data.

### Supplementary Table legends

Table S1: markers of major cell types calculated by FindVariableGenes function.

Table S2: Gene modules of epithelial root branch calculated by monocle3.

Table S3: Gene modules of SI branch calculated by monocle3.

Table S4: Gene modules of ameloblast branch calculated by monocle3.

Table S5: Enrichment of root branch gene module in transcription factor targets predicted by SCENIC.

Table S6: Enrichment of SI branch gene module in transcription factor targets predicted by SCENIC.

Table S7: Enrichment of ameloblast branch gene module in transcription factor targets predicted by SCENIC.

Table S8: Gene modules of mesenchyme development calculated by monocle3.

Table S9: Enrichment of mesenchyme development gene module in transcription factor targets predicted by SCENIC.

Table S10: ligand-target predictions of epithelial branch genes by nichenetr.

Table S11: ligand importance predictions of epithelial branch genes by nichenetr.

Table S12: ligand-target predictions of mesenchyme development genes by nichenetr.

Table S13: ligand importance predictions of mesenchyme development genes by nichenetr.
